# Pallidal Spectral and Phase-Amplitude Coupling Differences in Parkinson’s Disease Locomotor States

**DOI:** 10.64898/2025.12.28.696435

**Authors:** Josephine J. Wallner, Nicholas Druck, Harsh P. Shah, Leslie J. Cloud, Kathryn L. Holloway, Dean J. Krusienski

## Abstract

**Background:** Deep brain stimulation of the subthalamic nucleus or Globus Pallidus internus is an established therapy for Parkinson’s Disease that is sub-optimally managed with medication. While post-operative motor studies have defined neurophysiological signatures of the subthalamic nucleic local field potentials, comparable studies in the Globus pallidus internus are needed. Recent work shows that bandpower and phase-amplitude coupling in the subthalamic nucleus reliably distinguish locomotor states, motivating parallel investigation in the Globus pallidus internus.

**Objectives:** To characterize pallidal bandpower and phase-amplitude coupling across locomotor states (sitting, standing, and overground walking) in Parkinson’s, and relate to clinical motor scores.

**Methods:** Bandpower and phase-amplitude coupling from pallidal signals of six adults with Parkinson’s were compared across locomotor states; inter-state differences were correlated with clinical scores.

**Results:** High beta and gamma power differentiated sitting from walking, while delta power distinguished standing from walking. Phase-amplitude coupling between lower phases and gamma amplitude decreased during walking. Standing-to-walking delta-power modulation strongly correlated with Unified Parkinson’s Disease Rating Scale scores (R^2^ = 0.82, q < 0.05). Standing-to-walking beta-gamma phase-amplitude coupling modulation highly correlated with freezing-of-gait scores (R^2^ = 0.96, p < 0.05).

**Conclusions:** This study provides the first detailed characterization of Globus pallidus internus activity in these locomotor states in Parkinson’s. The coupling between lower frequencies and gamma was lower during walking compared to sitting and standing, revealing an inverse relationship to what has been reported in the subthalamic nucleus. These findings suggest that movement coordination in the Globus pallidus internus may rely on a balance of single- and cross-frequency mechanisms.

## 1 Introduction

Parkinson’s disease (PD) is a progressive neurodegenerative disorder characterized by bradykinesia, rigidity, tremor, and gait impairment [1]. When motor symptoms become inadequately controlled by dopaminergic medication, deep-brain stimulation (DBS) of the subthalamic nucleus (STN) or globus pallidus internus (GPi) offers an effective therapeutic alternative. While STN remains the predominant target, GPi-DBS has gained increasing clinical interest due to its favorable effects on levodopa-induced dyskinesias and a potentially more straightforward programming profile [2].

Advances in DBS technology now permit chronic recording of local field potentials (LFPs) from implanted electrodes, providing a window into the oscillatory dynamics of basal ganglia circuits. Extensive work in the STN has identified pathological beta-band (13–30 Hz) synchronization as a robust electrophysiological marker of the Parkinsonian state, with beta power correlating with bradykinesia and rigidity severity [3]. Beyond single-frequency measures, cross-frequency interactions, particularly phase-amplitude coupling (PAC) between beta phase and gamma amplitude, have emerged as candidate biomarkers. Beta-gamma PAC has been shown to reflect cortico-basal ganglia communication, motor network dysfunction, and changes in gait patterns [3–11].

These neurophysiological signatures have motivated the development of adaptive DBS (aDBS), which modulates stimulation delivery using real-time LFP features [12, 13]. The recent FDA approval of aDBS systems has accelerated efforts to identify biomarkers suitable for closed-loop control. In the STN, beta power during naturalistic locomotion has shown promise as an aDBS control signal [14]. However, analogous GPi biomarkers for aDBS remain underexplored.

The GPi serves as the primary output nucleus of the basal ganglia, integrating signals from the direct and indirect pathways before projecting to the thalamus and motor cortex. Prior work suggests that the STN and GPi exhibit reciprocal oscillatory patterns, with distinct and opposing modulation across these structures [15, 16]. Early intraoperative studies demonstrated that pallidal beta power and PAC with high-frequency oscillations were modulated during upper-limb movement [17]. Whether similar modulation occurs during whole-body locomotion, and whether such signatures relate to clinical motor scores, have not been extensively examined in the GPi.

In this study, we recorded GPi LFPs from six individuals with PD during a post-operative overground walking protocol. We characterized spectral power and PAC across three locomotor states: sitting, standing, and walking, and examined correlations with clinical motor assessments. Our objectives were to (1) identify GPi electrophysiological signatures that distinguish locomotor states, (2) compare these patterns to established STN findings, and (3) evaluate the potential of GPi spectral features as biomarkers for clinical motor dysfunction.

## 2 Methods

### 2.1 Participants and DBS devices

Six participants diagnosed with PD were recruited for the study. All participants reported persistent FoG following DBS implantation. The study design was approved by the Institutional Review Board of Virginia Commonwealth University, and informed consent was obtained for experimentation with human subjects. All participants were implanted with the Percept (Medtronic, Minneapolis, MN) DBS system in the GPi (unilateral or bilateral, based on clinical indication - see table in Figure S1) [18]. Each electrode shaft consists of four contacts, forming three bipolar recording channels (i.e., C0-C2, C1-C3, C0-C3). The Percept system records LFPs digitized at 250 Hz. All data was recorded using the Percept’s Indefinite Streaming mode, which is done with stimulation off. Patient demographics, clinical characteristics, and device setup can be found in Figure S1.

### 2.2 Data Collection

The participants walked an obstacle course that was originally designed to elicit FoG. Minimal freezing data were obtained during the experimental sessions and excluded from the subsequent analysis. The course consisted of five scenarios while LFPs were recorded: (1) passing a narrow archway, (2) 540^⍰^ turn, (3) approaching a visual target, (4) dual-task (performing calculations while walking), and (5) a time-sensitive task (answering a distant ringing phone). Data were collected off medication and with stimulation off. A summary of data length across participants is listed in Table 1.

**Table 1:** Seconds of data collected in each locomotor state across participants A–F.

| Participant \ Condition | A | B | C | D | E | F |
| --- | --- | --- | --- | --- | --- | --- |
| Sit | 120 | 127 | 10 | 38 | 60 | 102 |
| Stand | 120 | 12 | 120 | 30 | 51 | 20 |
| Walk | 120 | 130 | 130 | 120 | 98 | 120 |

During the experimental sessions, participants donned ankle-mounted inertial measurement units (IMUs), which were temporally aligned with GPi LFP recordings through the injection of a controlled external artifact. Transcutaneous electrical nerve stimulation (TENS) electrodes were positioned on the lateral neck, over the subcutaneous DBS leads, and on the contralateral mastoid. Surface EMG electrodes placed adjacent to the DBS leads detected the TENS artifact. Both EMG and IMU data streams were routed to a laptop via Lab Streaming Layer (LSL) for synchronized acquisition [19]. The TENS artifact, present in both the IMU and LFP signals, enabled offline temporal alignment.

Video recordings were acquired concurrently and synchronized to IMU and LFP data using timestamps and audio markers. Additionally, an external audio signal was transmitted through LSL to serve as a secondary synchronization marker across the video and IMU recordings. Based on the synchronized IMU and video data, the LFP recordings were manually segmented into three locomotor states: sitting, standing, and walking. Prior to LFP data collection, each participant underwent motor testing: Unified Parkinson’s Disease Rating Scale section III (UPDRS-III) and the new freezing-of-gait questionnaire (nFoG-Q).

### 2.3 Data Analysis

#### 2.3.1 Preprocessing

To provide a balanced comparison across participants with differing numbers of implants, a single hemisphere was chosen for analysis. For those with bilateral implants, GPi contacts contralateral to hemibody with worse (ie., greater) UPDRS-III scores at the time of collection were chosen. One participant had equal UPDRS-III scores bilaterally; for this participant, the initially affected hemisphere per prior clinical documentation was used. For all participants, the C1 and C2 contacts were confirmed to be located in the GPi by a neurosurgeon via post-operative imaging using Stealth MRI (Medtronic, Minneapolis, MN) [20]. From these contacts, two bipolar channels based on the C0-C2 and C1-C3 contact pairs were used for the subsequent analyses. Channel C0-C3 was not analyzed due to the lower spatial specificity and overlap with the selected channels. The raw LFP data and power spectra were screened to confirm the absence of motion artifacts and for EKG contamination using the Perceive toolbox [21]. Dual-tasking walking data was excluded to avoid confounding effects in the signals. While some participants had shorter lengths of clean LFPs, the subsequent analysis techniques have been shown to be robust for these segment lengths [22].

#### 2.3.2 Frequency Bandpower

Bandpower estimates were calculated to quantify the relative contributions of physiological frequencies across locomotor states. The power spectral densities (PSDs) were estimated for each locomotor state using Welch’s method via the *pwelch* function in MATLAB2024a (MathWorks Inc., Natick, MA). The LFPs were partitioned into 1-second Hamming-windowed epochs with 50% overlap. Due to the lowpass filter characteristics of the Percept, the resulting PSDs were processed and visualized up to 100 Hz.

To reveal within-participant differences in bandpower between locomotor states, the *bandpower* function in MATLAB2024a was used across canonical frequency bands (i.e., Delta: 0.5-4 Hz, Theta: 4-8 Hz, Alpha: 8-12 Hz, Beta: 13-30 Hz, and Gamma: 30-100 Hz). Based on prior work suggesting functionally relevant subdivisions of the beta band in the GPi, low beta (13-20 Hz) and high beta (21-30 Hz) were also analyzed [17].

Pairwise statistical comparisons between locomotor states were performed separately for each frequency band using the Wilcoxon signed-rank test, a nonparametric test that does not assume normality. P-values were adjusted for multiple comparisons using Benjamini-Hochberg correction (significance threshold: 0.05).

#### 2.3.3 Phase-Amplitude Coupling

Phase–Amplitude Coupling (PAC) is a measure of cross-frequency coupling where the phase of a low frequency oscillation modulates the amplitude of a higher-frequency oscillation. PAC was quantified using the modulation index (MI) based on Kullback–Leibler divergence, a widely-used method in LFP studies [17, 22, 23]. LFP signals were bandpass filtered to estimate phase frequencies from 1–30 Hz (1 Hz steps, 2 Hz bandwidth) and amplitude frequencies from 40–100 Hz (2 Hz steps, 10 Hz bandwidth). Instantaneous phase and amplitude were extracted using the Hilbert transform. Specifically, for every 20-degree interval of the instantaneous phase distribution, the entropy of the instantaneous amplitude envelope distribution was computed.

MI values were computed between each phase–amplitude frequency pair and visualized as comodulograms. To obtain summary PAC metrics, MI values were summed across canonical phase-frequency ranges, consistent with prior DBS work [5]:

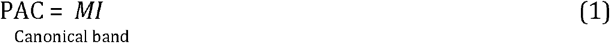

Where *canonical band* represents the canonical frequency band used to define the phase frequency range for the summation of MI values. This process was repeated for each locomotor state.

Pairwise statistical comparisons between locomotor states were performed using Wilcoxon signed rank tests. Correction for multiple comparisons was done using Benjamini–Hochberg FDR within each spectral feature family separately (band power and PAC). A significance threshold of 0.05 was used.

#### 2.3.4 Clinical Correlations

Spectral features that showed significant state-dependent modulation were averaged across the two GPi contacts within each patient, and tested against two clinical scores (UPDRS-III and nFoG-Q) using ordinary least-squares linear regression. Regression diagnostics were inspected for each fit, and outliers were identified using the Bonferroni-corrected outlier test on externally studentized residuals; where flagged, regressions were refit with that observation excluded, and both full-cohort and outlier-excluded results are reported. Benjamini-Hochberg FDR correction was applied within each spectral-metric × clinical-scale family. Band-power and PAC correlations were treated as separate families, as they test distinct neurophysiological mechanisms (oscillatory power vs cross-frequency coupling). All statistical tests were two-tailed.

## 3 Data Sharing

Data are available upon request.

## 4 Results

### 4.1 Differences in frequency bandpower across locomotor states

Figure 1a shows across-participant average for all three locomotor states: sit, stand, and walk. To assess statistically significant, individual within-participant bandpower differences across locomotor states, the bandpower in each canonical frequency band was computed across each participant’s GPi contacts for each state, and displayed as box plots in Figure 1b. Individual PSD for all GPi contacts as shown in Figure S2.

**Figure 1:**
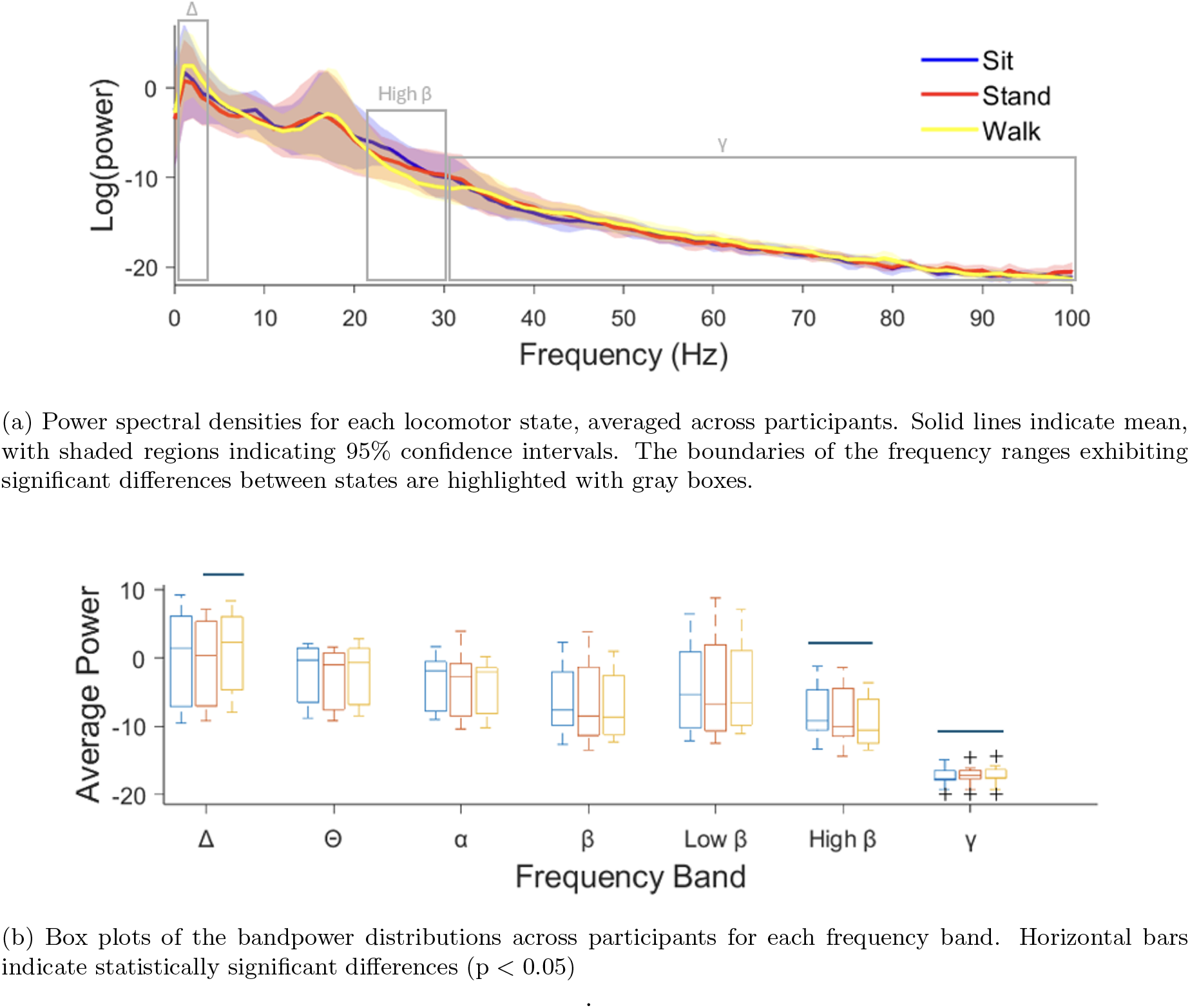
Summary of bandpower analysis across locomotor states.

Pairwise comparisons using the Wilcoxon signed-rank test with Benjamin-Hochberg correction between locomotor states revealed increased delta power during walking relative to standing. Power in both high beta and gamma frequency ranges differentiated between sitting and walking: walking was associated with decreased high beta and increased gamma. Notably, the beta power modulation between sitting and walking was not apparent when analyzing the entire beta range as a single band.

### 4.2 Differences in phase-amplitude coupling across locomotor states

Figure 2a provides example comodulograms for one contact from a selected participant across locomotor states. The insets highlight the coupling between the phase of canonical frequency ranges (i.e., theta, alpha, beta) and the gamma amplitude. PAC values from these comodulograms were summarized into locomotor-state-specific boxplots, as shown in Figure 2b. Individual comodulograms for GPi contacts across all participants are shown in Figure S3.

**Figure 2:**
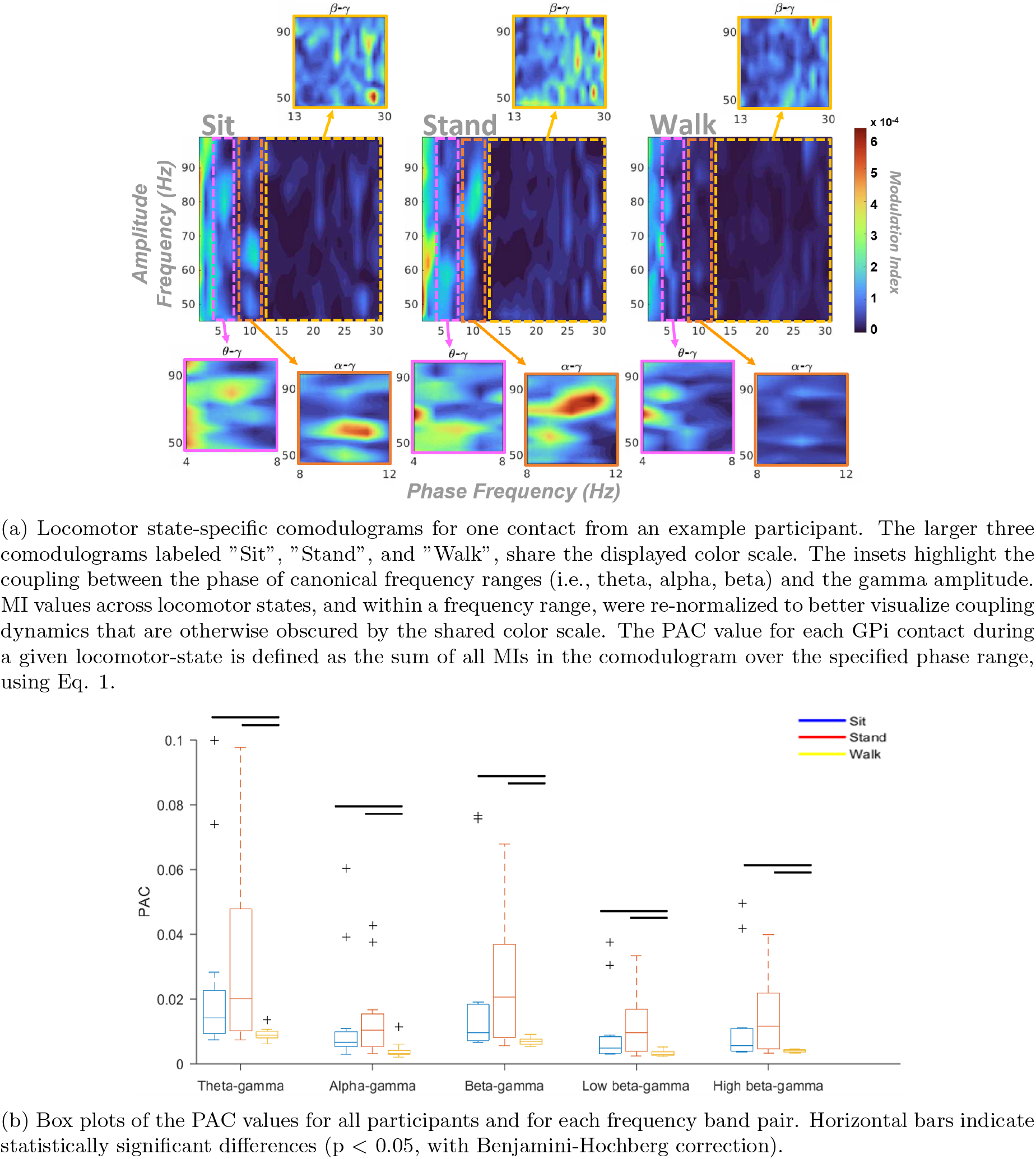
Summary of the current GPi PAC results.

No significant differences in PAC were observed between sitting and standing. However, compared to sitting and standing, walking was consistently associated with a decrease in PAC between all lower frequency bands and gamma. Standing-related PAC is highest at low phase frequencies and gradually declines with increasing phase frequency, similar to the typical 1/f-like decay in neural power spectra; however, these PAC patterns cannot be fully accounted for by frequency-dependent changes in power [24]. This difference across locomotor states was present whether or not the beta band was partitioned into low and high beta; therefore, subsequent PAC comparisons were done using the entire beta band without subdivisions.

Figure 3 compares the current GPi beta–gamma PAC results with previously published STN findings from Farokhniaee et al. (2024) [5]. To enable an analogous comparison, the beta–gamma PAC values from Figure 2b were min–max normalized to match the scale in the STN analysis. In this STN study, beta-gamma PAC was positively modulated during walking across their participants; whereas in this GPi cohort, a reciprocal trend was observed where beta-gamma PAC was negatively modulated during walking.

**Figure 3:**
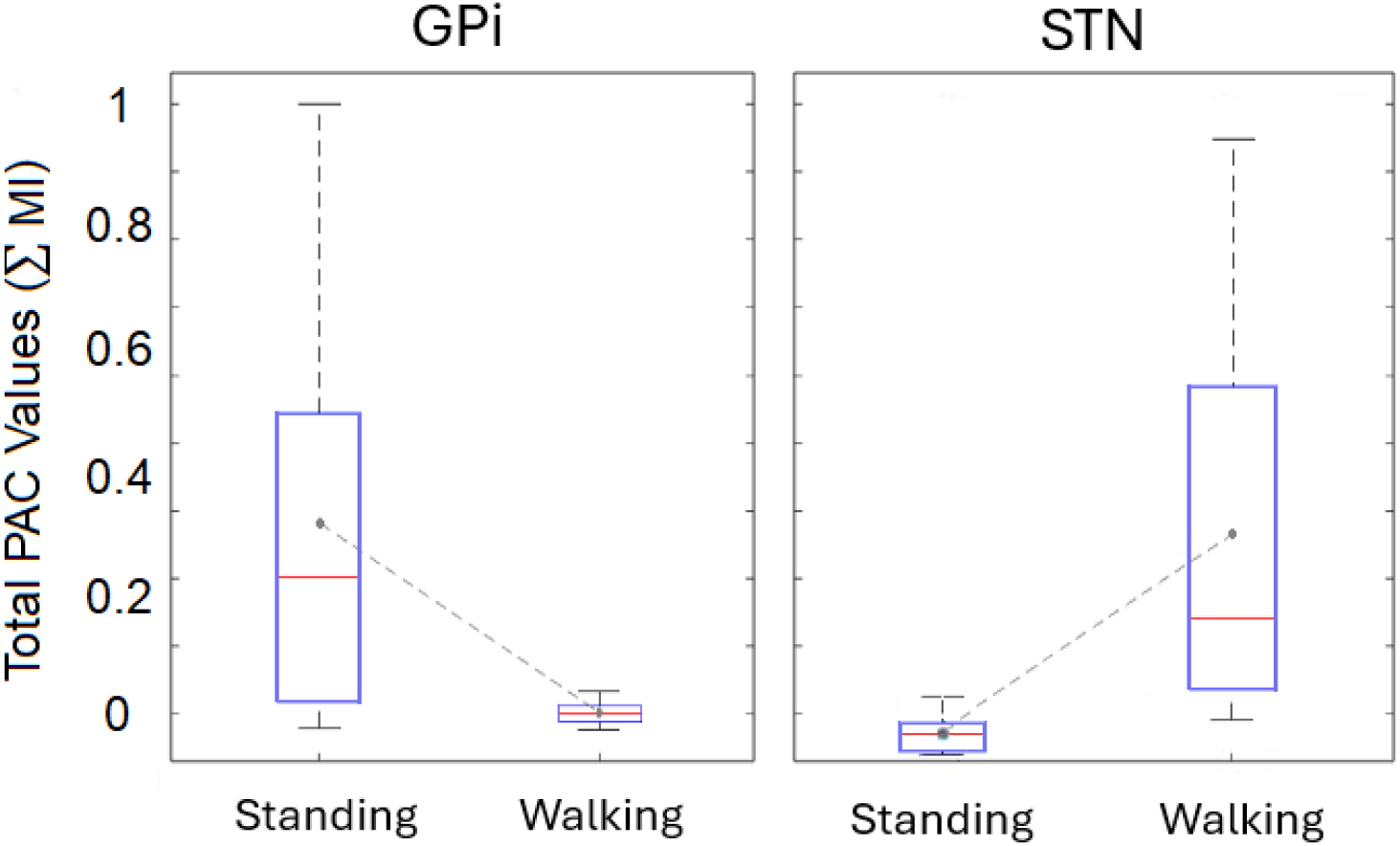
Min-max normalized changes in beta-gamma PAC between standing and walking, compared between the present GPi participants and a separate STN cohort from Farokhniaee et al., 2024. Beta-gamma PAC between standing and walking exhibits a reciprocal relationship across STN and GPi.

#### 4.3 Correlations between spectral features and clinical motor scores

Figure 4 illustrates the results of the clinical correlations. δ band power Stand⍰Walk modulation correlated positively with MDS-UPDRS-III scores (R^2^ = 0.82, slope β = +4.04, 95% CI [+1.40, +6.68]; q = 0.040; n = 6, no outliers flagged). β-γ PAC Stand⍰Walk modulation correlated with nFOG-Q scores after outlier exclusion of subject 0411 (flagged by the Bonferroni-corrected outlier test): R^2^ = 0.96, slope β = +159.9, 95% CI [+103.4, +216.5]; q = 0.007; n = 5. The full-cohort regression (n = 6) was not significant. Given the cohort size, the clinical correlations are reported as exploratory pending replication.

**Figure 4:**
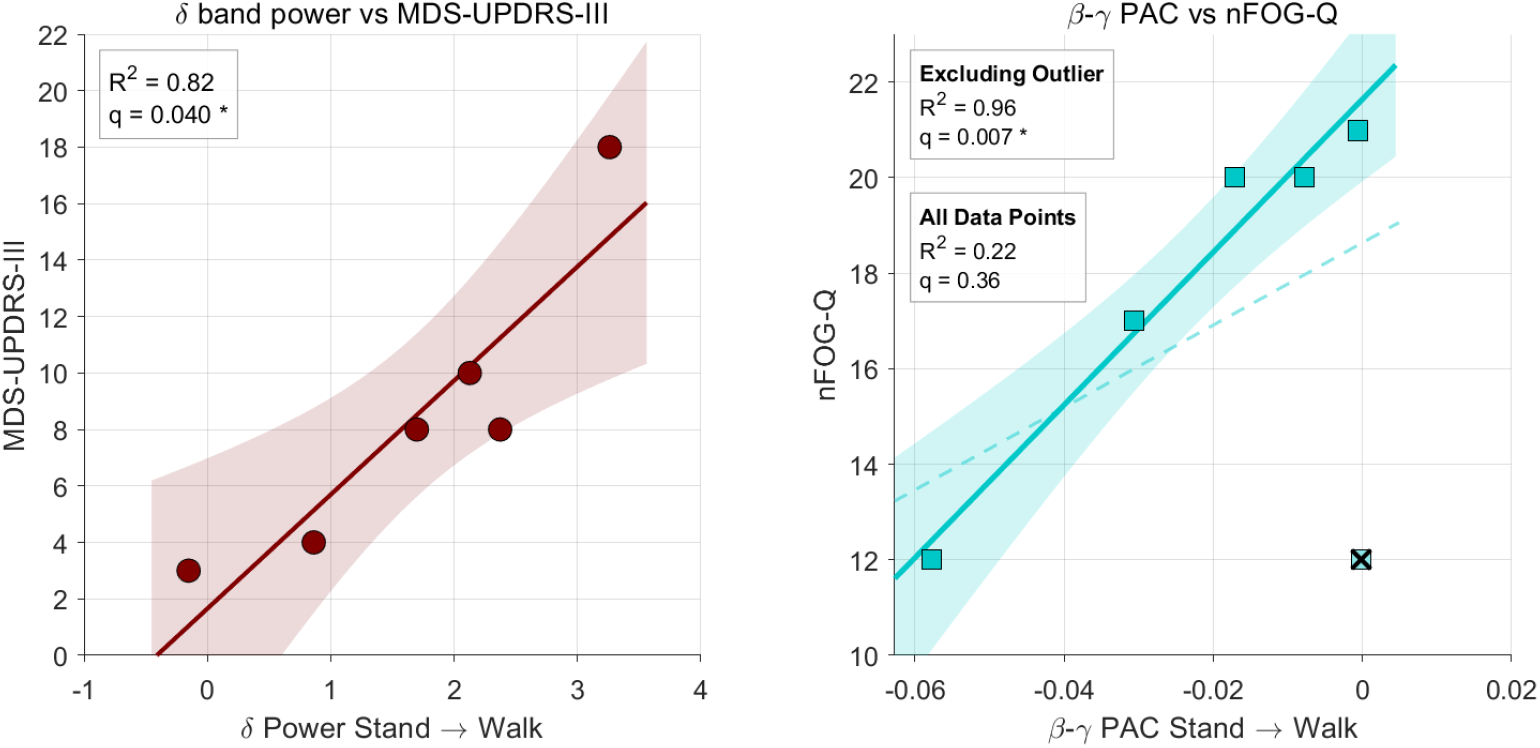
Left: Stand→Walk increase in *δ* band power vs MDS-UPDRS-III scores (n = 6). Right: Stand→Walk modulation of *β*-*γ* PAC vs nFOG-Q scores. Dashed line and faded points, full cohort (n = 6); solid line and shaded 95% confidence band, outlier-excluded fit (n = 5). The outlier participant (×) was flagged by the Bonferroni-corrected outlier test on externally studentized residuals. R^2^ and BH-FDR-adjusted q reported within each panel; * indicates q *<* 0.05.

## 5 Discussion

The present study, to our knowledge, provides the first detailed characterization of GPi LFP dynamics during naturalistic sitting, standing, and overground walking in individuals with PD. High-beta and gamma power differentiated sitting from walking, while delta power distinguished standing from walking. PAC between lower-frequency phases (theta, alpha, beta) and gamma amplitude was consistently reduced during walking compared to static states. Standing-to-walking delta modulation correlated strongly with motor scores (UPDRS-III), and beta-gamma PAC modulation tracked with freezing-of-gait scores (nFoG-Q).

The observed reduction in high-beta power during walking aligns with the well-established phenomenon of movement-related beta desynchronization, which has been extensively documented in the STN and is considered a hallmark of healthy motor output [25, 26]. Notably, this effect was only apparent when analyzing high-beta separately from low-beta, suggesting that beta sub-bands may reflect distinct functional processes in the GPi, as previously proposed for the STN [27, 28].

The increased activity in the 1-8 Hz range during walking, which has also been reported in the pedunculopontine nucleus, suggests that low-frequency oscillations may be a shared gait-related signature across interconnected nodes of the motor network [29]. The accompanying increase in gamma power during walking further indicates that multiple oscillatory components reconfigure as participants transition between static and dynamic states, underscoring the distributed, multi-frequency nature of motor control in basal ganglia circuits.

A key finding was the global reduction in PAC between all lower-frequency phases and gamma amplitude during walking. This pattern, where sit ≈ stand > walk, suggests that cross-frequency coupling across a broad frequency range in the GPi may encode a movement-related control signal. Cross-frequency coupling has been proposed to reflect dynamic gating mechanisms that regulate when neural populations become synchronized in response to behavioral context [30, 31]. The observed reduction during active movement could reflect the release of an inhibitory signal from the GPi, facilitating motor execution.

Notably, the direction of beta-gamma PAC modulation in this GPi cohort was opposite to that reported in the STN, where PAC increases during walking [5]. This reciprocal pattern is consistent with previous work in the STN and GPi, which reported inverse oscillatory dynamics in response to therapeutic interventions [15, 32]. While the present study did not include simultaneous STN recordings, future work employing dual-target acquisition could directly test this reciprocal relationship during locomotion.

The identification of locomotor-state-dependent spectral features in the GPi carries implications for aDBS development. Incorporating multi-band, state-aware biomarkers, such as delta modulation or beta-gamma PAC, may enhance adaptive control by accounting for the specific motor context. Furthermore, longitudinal tracking of these markers could provide physiological indices of disease progression or short-timescale fluctuations in motor state, potentially guiding individualized parameter adjustments over time.

### 5.1 Limitations

The sample size (n = 6) restricts generalizability and statistical power, particularly for the clinical correlation analyses, which should be interpreted as hypothesis-generating in the absence of a larger cohort. Participants exhibited variable disease duration and medication regimens, which may have influenced basal ganglia activity. Additionally, while this study was completed with DBS OFF, the slow dissipative effects of previous DBS stimulation when DBS was ON in the GPi likely contributed to inter-subject variability [33]. Finally, while PAC has shown discriminatory value across locomotor states, the observed broadband coupling suggests that PAC may not be optimally identifying processes that drive these differences.

### 5.2 Summary and conclusion

This study demonstrates that GPi LFPs encode locomotor-state-dependent information that can be captured by single-frequency power and cross-frequency coupling measures. The reciprocal PAC modulation relative to the STN suggests that movement coordination in the pallidum may involve distinct, and potentially reciprocal, oscillatory dynamics. The observed correlations with clinical motor scores, while preliminary, support continued investigation of GPi spectral features as candidate biomarkers for pallidal aDBS and as potential indices of motor dysfunction in PD.

## 6 Acknowledgments

This work was funded in part by 2022 & 2024 Virginia Commonwealth University Parkinson’s and Movement Disorders Center (PMDC) Pilot Grants. The authors would also like to thank Gina Marie Blackwell for her assistance in obtaining the IRB approval and patient coordination; Alex Feria for data collection; and Ken Koltermann and Gang Zhou for providing the ankle inertial measurement units and their associated synchronization software.

## 7 Authors’ Roles

Conceptualization and study design: JW, HS, LC, KH, DJK; Data collection: JW, ND, HS, LC, KH; Formal analysis: JW, ND; Manuscript preparation: JW, ND, DJK; Manuscript review & editing: JW, ND, HS, LC, KH, DJK

## References

[1] Saman Zafar and Sridhara S Yaddanapudi. Parkinson disease. In StatPearls. StatPearls Publishing, Treasure Island (FL), January 2025.

[2] Shi-Ying Fan, Kai-Liang Wang, Wei Hu, Robert S Eisinger, Alexander Han, Chun-Lei Han, Qiao Wang, Shimabukuro Michitomo, Jian-Guo Zhang, Feng Wang, Adolfo Ramirez-Zamora, and FanGang Meng. Pallidal versus subthalamic nucleus deep brain stimulation for levodopa-induced dyskinesia. Ann. Clin. Transl. Neurol., 7(1):59–68, January 2020.

[3] Thomas Koeglsperger, Jan H Mehrkens, and Kai Bötzel. Bilateral double beta peaks in a PD patient with STN electrodes. Acta Neurochir. (Wien), 163(1):205–209, January 2021.

[4] Natasha Darcy, Roxanne Lofredi, Bassam Al-Fatly, Wolf-Julian Neumann, Julius Hübl, Christof Brücke, Patricia Krause, Gerd-Helge Schneider, and Andrea Kühn. Spectral and spatial distribution of subthalamic beta peak activity in parkinson’s disease patients. Exp. Neurol., 356(114150):114150, October 2022.

[5] Amirali Farokhniaee, Chiara Palmisano, Jasmin Del Vecchio Del Vecchio, Gianni Pezzoli, Jens Volkmann, and Ioannis U Isaias. Gait-related beta-gamma phase amplitude coupling in the subthalamic nucleus of parkinsonian patients. Sci. Rep., 14(1):6674, March 2024.

[6] Andrew I Yang, Nora Vanegas, Codrin Lungu, and Kareem A Zaghloul. Beta-coupled highfrequency activity and beta-locked neuronal spiking in the subthalamic nucleus of parkinson’s disease. J. Neurosci., 34(38):12816–12827, September 2014.

[7] Tisa Hodnik, Stiven Roytman, Nico I Bohnen, and Uros Marusic. Beta-gamma phase-amplitude coupling as a non-invasive biomarker for parkinson’s disease: Insights from electroencephalography studies. Life (Basel), 14(3):391, March 2024.

[8] Yuki Kimoto, Naoki Tani, Takuto Emura, Takahiro Matsuhashi, Takuto Yamamoto, Yuya Fujita, Satoru Oshino, Koichi Hosomi, Hui Ming Khoo, Shimpei Miura, Takahiro Fujinaga, Takufumi Yanagisawa, and Haruhiko Kishima. Beta-gamma phase-amplitude coupling of scalp electroencephalography during walking preparation in parkinson’s disease differs depending on the freezing of gait. Front. Hum. Neurosci., 18:1495272, November 2024.

[9] Haifeng Zhao, Shenglin Hao, Peiran Zhang, Shenghong He, Laura Wehmeyer, Ziyi Feng, Lu Xu, Shikun Zhan, Wei Liu, Xiaoxiao Zhang, Marie-Laure Welter, Dianyou Li, Bomin Sun, Yong Lu, Huiling Tan, and Chunyan Cao. Cortical and corticomuscular beta-gamma phase-amplitude coupling during different locomotion states and the effects of levodopa in parkinson’s disease. Mov. Disord., (mds.70031), September 2025.

[10] Philipp A. Loehrer, Sahar Yassine, Immo Weber, Valentin Sanner, Shenghong He, Alek Pogosyan, Lijiao Chen, Laura Witt, Gereon Rudolf Fink, David J. Pedrosa, Lars Timmermann, and Huiling Tan. Characterizing the frequency-specific and spatiotemporal dynamics of - phase-amplitude coupling in parkinson’s disease. bioRxiv, 2025.

[11] Zhen Yin, Guoping Zhu, Yuchen Liu, Bin Zhao, Dandan Liu, Yiqing Bai, Qiqi Zhang, Lei Shi, Tao Feng, Aiping Yang, Hongli Liu, Fangang Meng, Wolfgang J Neumann, Andrea A Kühn, Yuan Jiang, and Jianfeng Zhang. Cortical phase-amplitude coupling is key to the occurrence and treatment of freezing of gait. Brain, 145(7):2407–2421, 2022.

[12] Simon Little, Alex Pogosyan, Spencer Neal, Baltazar Zavala, Ludvic Zrinzo, Marwan Hariz, Thomas Foltynie, Patricia Limousin, Keyoumars Ashkan, James FitzGerald, Alexander L Green, Tipu Z Aziz, and Peter Brown. Adaptive deep brain stimulation in advanced parkinson disease. Ann. Neurol., 74(3):449–457, September 2013.

[13] Kevin B Wilkins, Matthew N Petrucci, Emilia F Lambert, Jillian A Melbourne, Aryaman S Gala, Pranav Akella, Laura Parisi, Chuyi Cui, Yasmine M Kehnemouyi, Shannon L Hoffman, Sudeep Aditham, Cameron Diep, Hannah J Dorris, Jordan E Parker, Jeffrey A Herron, and Helen M Bronte-Stewart. Beta burst-driven adaptive deep brain stimulation for gait impairment and freezing of gait in parkinson’s disease. Brain Commun., 7(4):fcaf266, July 2025.

[14] Shenghong He, Fahd Baig, Anca Merla, Flavie Torrecillos, Andrea Perera, Christoph Wiest, Jean Debarros, Moaad Benjaber, Michael G Hart, Lucia Ricciardi, Francesca Morgante, Harutomo Hasegawa, Michael Samuel, Mark Edwards, Timothy Denison, Alek Pogosyan, Keyoumars Ashkan, Erlick Pereira, and Huiling Tan. Beta-triggered adaptive deep brain stimulation during reaching movement in parkinson’s disease. Brain, 146(12):5015–5030, December 2023.

[15] Christoph Wiest, Flavie Torrecillos, Alek Pogosyan, Manuel Bange, Muthuraman Muthuraman, Sergiu Groppa, Natasha Hulse, Harutomo Hasegawa, Keyoumars Ashkan, Fahd Baig, Francesca Morgante, Erlick A Pereira, Nicolas Mallet, Peter J Magill, Peter Brown, Andrew Sharott, and Huiling Tan. The aperiodic exponent of subthalamic field potentials reflects excitation/inhibition balance in parkinsonism. eLife, 12:e82467, feb 2023.

[16] Stephen L. Schmidt, David T. Brocker, Brandon D. Swan, Dennis A. Turner, and Warren M. Grill. Evoked potentials reveal neural circuits engaged by human deep brain stimulation. Brain Stimulation, 13(6):1706–1718, 2020.

[17] Nicholas AuYong, Mahsa Malekmohammadi, Joni Ricks-Oddie, and Nader Pouratian. Movementmodulation of local power and phase amplitude coupling in bilateral globus pallidus interna in parkinson disease. Front. Hum. Neurosci., 12:270, July 2018.

[18] Medtronic, Inc. Percept PC deep brain stimulator, 2020. https://www.medtronic.com.

[19] Lsl-website. https://labstreaminglayer.org/#/. Accessed: 2025-11-14.

[20] Neurosurgery navigation stealthstation™ s8 surgical navigation system. http://web.archive.org/web/20080207010024/ http://www.808multimedia.com/winnt/kernel.htm.

[21] neuromodulation. perceive: MATLAB toolbox for extracting data from the medtronic percept bidirectional brain–computer interface. https://github.com/neuromodulation/perceive, 2020. Accessed: 2025-11-14.

[22] Adriano B L Tort, Robert Komorowski, Howard Eichenbaum, and Nancy Kopell. Measuring phase-amplitude coupling between neuronal oscillations of different frequencies. J. Neurophysiol., 104(2):1195–1210, August 2010.

[23] Jon López-Azćarate, Mikel Tainta, María C Rodríguez-Oroz, Miguel Valencia, Rafael González, Jorge Guridi, Jorge Iriarte, José A Obeso, Julio Artieda, and Manuel Alegre. Coupling between beta and high-frequency activity in the human subthalamic nucleus may be a pathophysiological mechanism in parkinson’s disease. J. Neurosci., 30(19):6667–6677, May 2010.

[24] Charmaine Demanuele, Christopher J James, and Edmund Js Sonuga-Barke. Distinguishing low frequency oscillations within the 1/f spectral behaviour of electromagnetic brain signals. Behav. Brain Funct., 3(1):62, December 2007.

[25] Clara Moisello, Daniella Blanco, Jing Lin, Priya Panday, Simon P Kelly, Angelo Quartarone, Alessandro Di Rocco, Chiara Cirelli, Giulio Tononi, and M Felice Ghilardi. Practice changes beta power at rest and its modulation during movement in healthy subjects but not in patients with parkinson’s disease. Brain Behav., 5(10):e00374, October 2015.

[26] Varvara Mathiopoulou, Roxanne Lofredi, Lucia K Feldmann, Jeroen Habets, Natasha Darcy, WolfJulian Neumann, Katharina Faust, Gerd-Helge Schneider, and Andrea A Kühn. Modulation of subthalamic beta oscillations by movement, dopamine, and deep brain stimulation in parkinson’s disease. NPJ Parkinsons Dis., 10(1):77, April 2024.

[27] Joohi Jimenez-Shahed, Ilknur Telkes, Ashwin Viswanathan, and Nuri F Ince. GPi oscillatory activity differentiates tics from the resting state, voluntary movements, and the unmedicated parkinsonian state. Front. Neurosci., 10:436, September 2016.

[28] Musa Ozturk, Aviva Abosch, David Francis, Jianping Wu, Joohi Jimenez-Shahed, and Nuri F Ince. Distinct subthalamic coupling in the on state describes motor performance in parkinson’s disease. Movement Disorders, 35(1):91–100, 2020.

[29] Rene Molina, Chris J Hass, Kristen Sowalsky, Abigail C Schmitt, Enrico Opri, Jaime A Roper, Daniel Martinez-Ramirez, Christopher W Hess, Kelly D Foote, Michael S Okun, and Aysegul Gunduz. Neurophysiological correlates of gait in the human basal ganglia and the PPN region in parkinson’s disease. Front. Hum. Neurosci., 14:194, June 2020.

[30] Xian Jiang, Jorge Gonzalez-Martinez, and Eric Halgren. Inter-area theta-gamma phase–amplitude coupling supports directed communication in the primate brain. Scientific Reports, 9:13487, 2019.

[31] Ilknur Telkes, Ashwin Viswanathan, Joohi Jimenez-Shahed, Aviva Abosch, Musa Ozturk, Akshay Gupte, Joseph Jankovic, and Nuri F Ince. Local field potentials of subthalamic nucleus contain electrophysiological footprints of motor subtypes of parkinson’s disease. Proceedings of the National Academy of Sciences, 115(36):E8567–E8576, 2018.

[32] Atsushi Nambu, Hironobu Tokuno, and Masahiko Takada. Functional significance of the cortico– subthalamo–pallidal ‘hyperdirect’ pathway. Neuroscience Research, 43(2):111–117, 2002.

[33] Scott E Cooper, Cameron C McIntyre, Hubert H Fernandez, and Jerrold L Vitek. Association of deep brain stimulation washout effects with parkinson disease duration. JAMA Neurol., 70(1):95– 99, January 2013.

