## Supplemental Material for "Pallidal Spectral and Phase-Amplitude Coupling Differences in Parkinson’s Disease Locomotor States"

***for***

***Clinical characteristics and DBS device***

Figure S1 compiles summary statistics for the current study participants across several demographic and clinical traits and provides an example schematic of the deep brain stimulation percept lead contacts.

***Power Spectral Densities (PSDs)***
The total PSDs for the middle two GPi-containing DBS contacts from the right and left hemispheres were computed for each patient (denoted by the data collection date) and for each locomotor state (sitting, standing, and walking). LFP signals from all moments of each state were concatenated and analyzed using 1-second windows with 50% overlap. PSDs were estimated, decibel-normalized, and plotted on a logarithmic scale. In Figure S2, PSDs are shown as thick lines with the following color scheme: blue for sitting, red for standing, and yellow for walking.

***Phase-Amplitude Coupling (PAC)***
Methodological details for PAC are provided in the main text (Results section). Individual phase-amplitude comodulograms were estimated for each GPi-containing contact across participants during sitting, standing, and walking. These comodulograms are presented in Figure S3.

***PAC Surrogate Testing***
To assess the statistical significance of PAC estimates, 500 surrogate comodulograms were generated for each GPi contact to create a null distribution of PAC values. Surrogates were generated by first computing the phase time series via the Hilbert transform, then randomly splitting the phase series into two segments, swapping their order, and recomputing PAC with the unaltered gamma amplitude envelope. For each phase-amplitude frequency pair, a proxy p-value was calculated as the proportion of surrogates with PAC values greater than or equal to the original PAC value. The incidence of GPi contacts with significant p-values (<0.05) at every phase-amplitude pair was compiled and visualized as “incidence plots” in Figure S4.

***Clinical Correlations***
Differences in bandpower and PAC were computed between each pair of locomotor states (i.e., Sit → Stand, Sit → Walk, Stand → Walk). Robust linear regression models were used to correlate absolute changes in both bandpower and PAC with clinical motor scores. Robust linear regression was used when there existed a need to address obvious outliers. Results of the regression analysis can be found in Figures S5A (bandpower) and S5B (PAC).

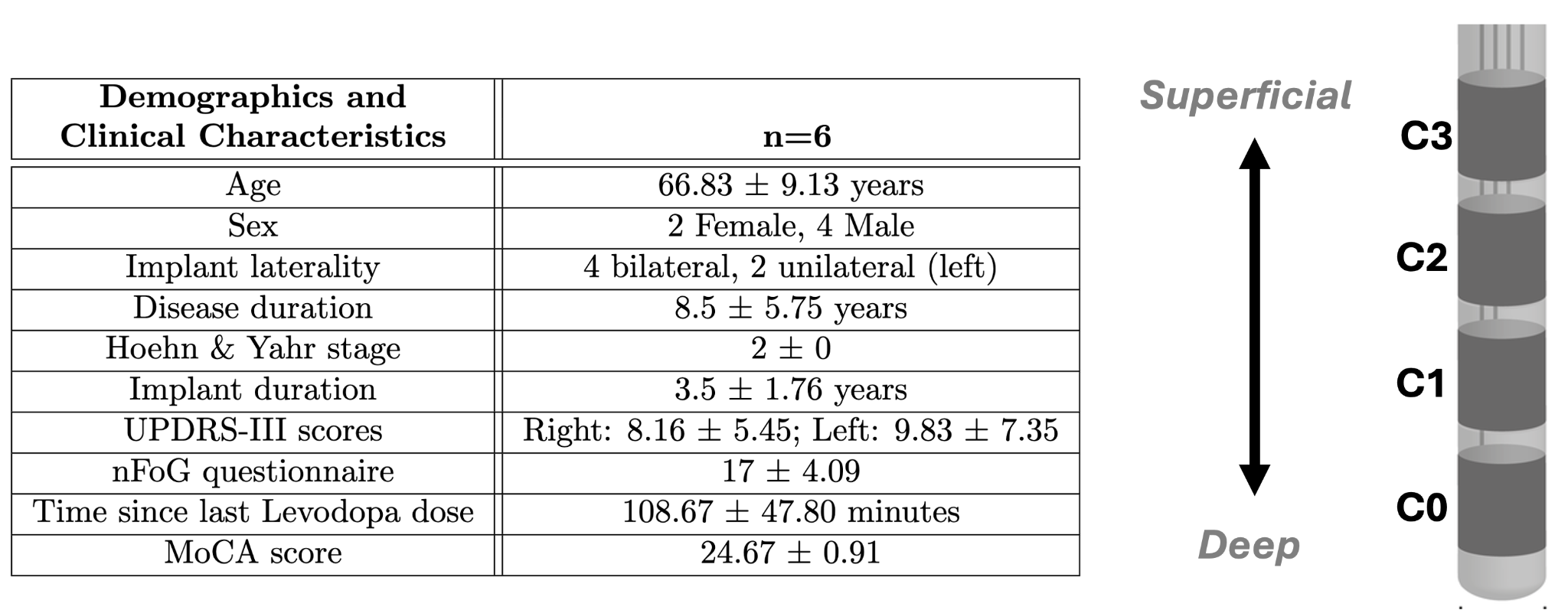

**Figure S1.** Participant demographics and clinical characteristics (left panel). Schematic of single DBS lead. C0-C3 represents individual contacts (right panel).

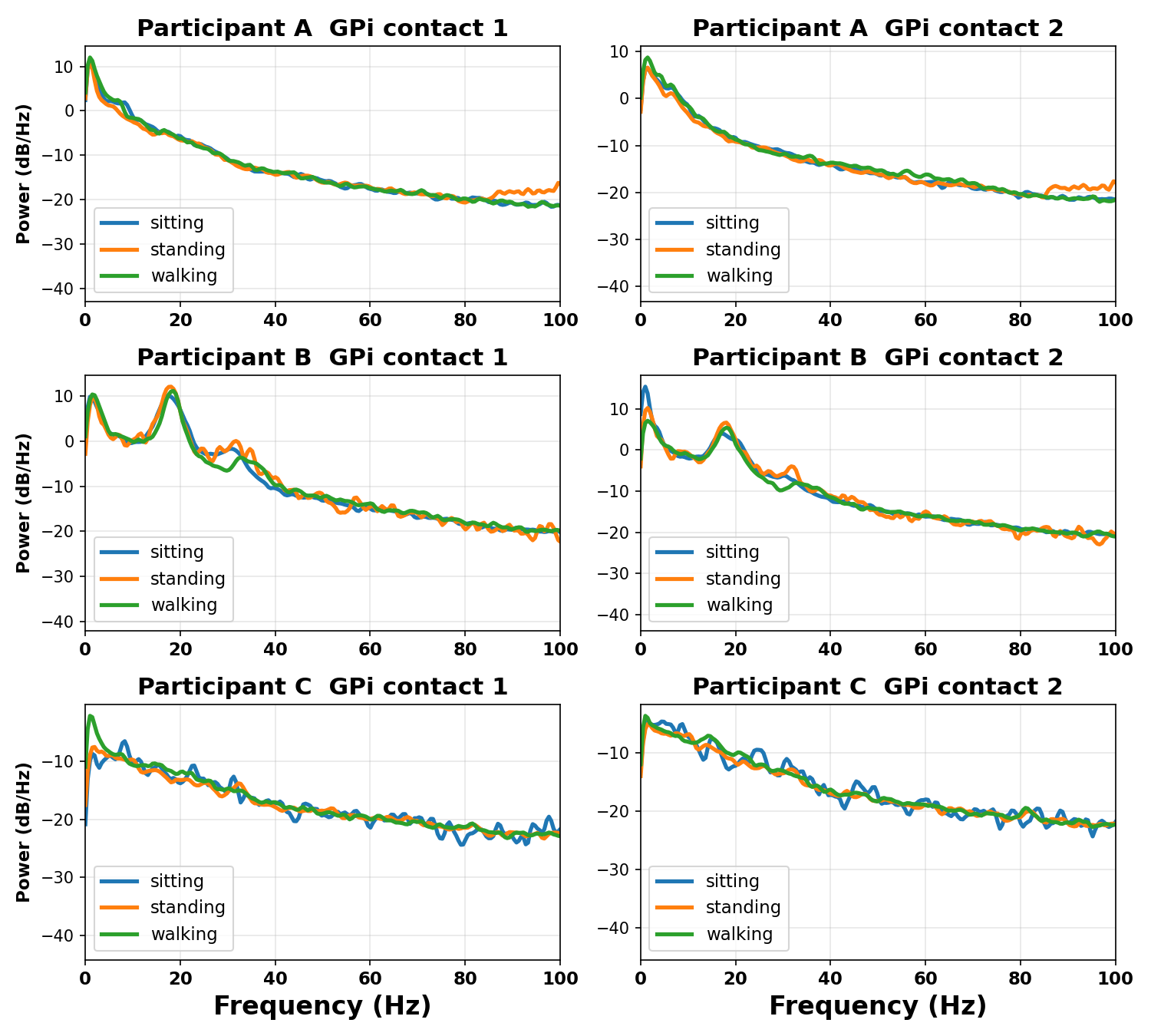

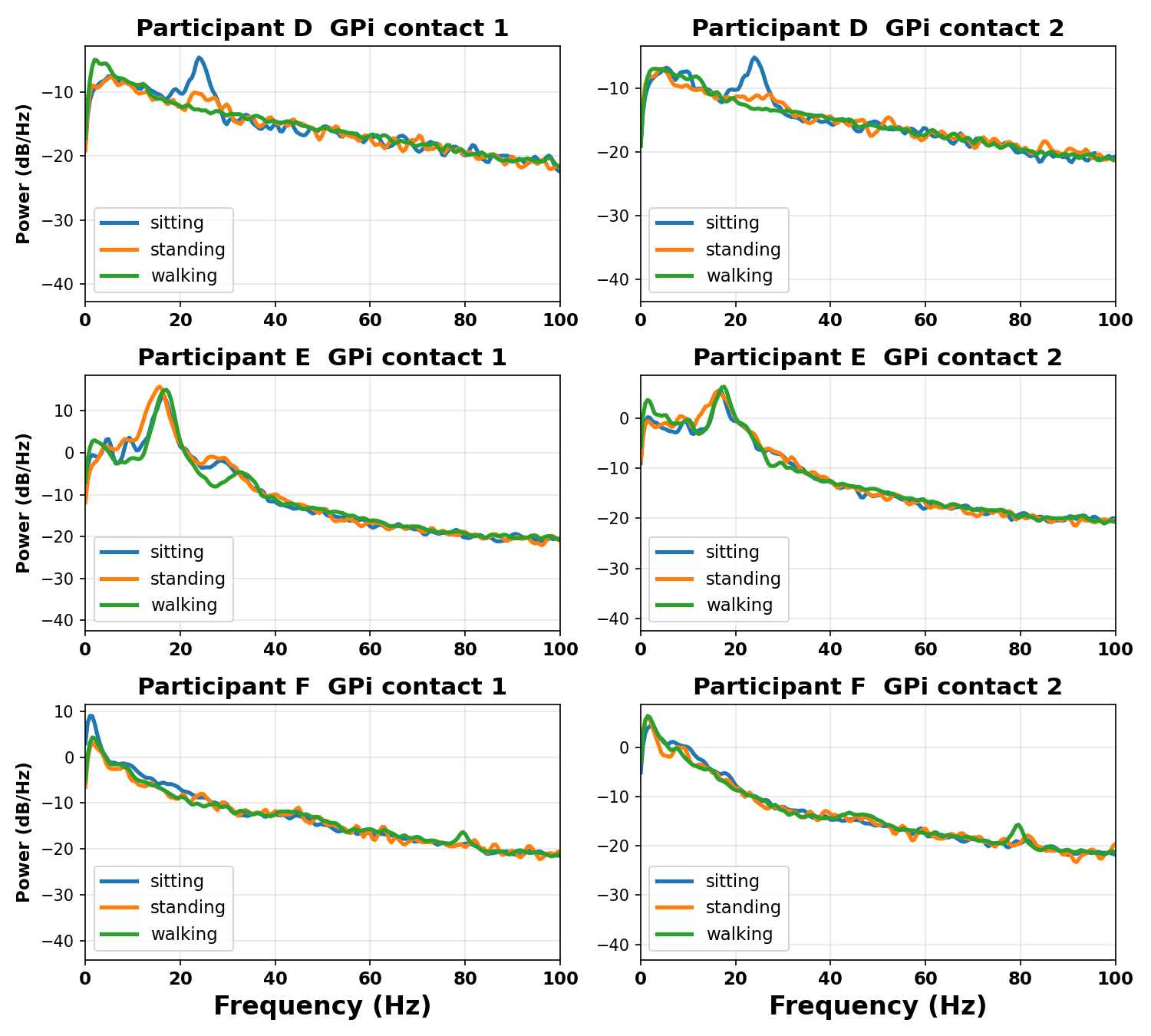

**Figure S2**. **Power Spectral Densities (PSDs).** Each subplot represents the PSDs for one of two GPi contacts used per participant across the three locomotor states (sit, stand, walk).

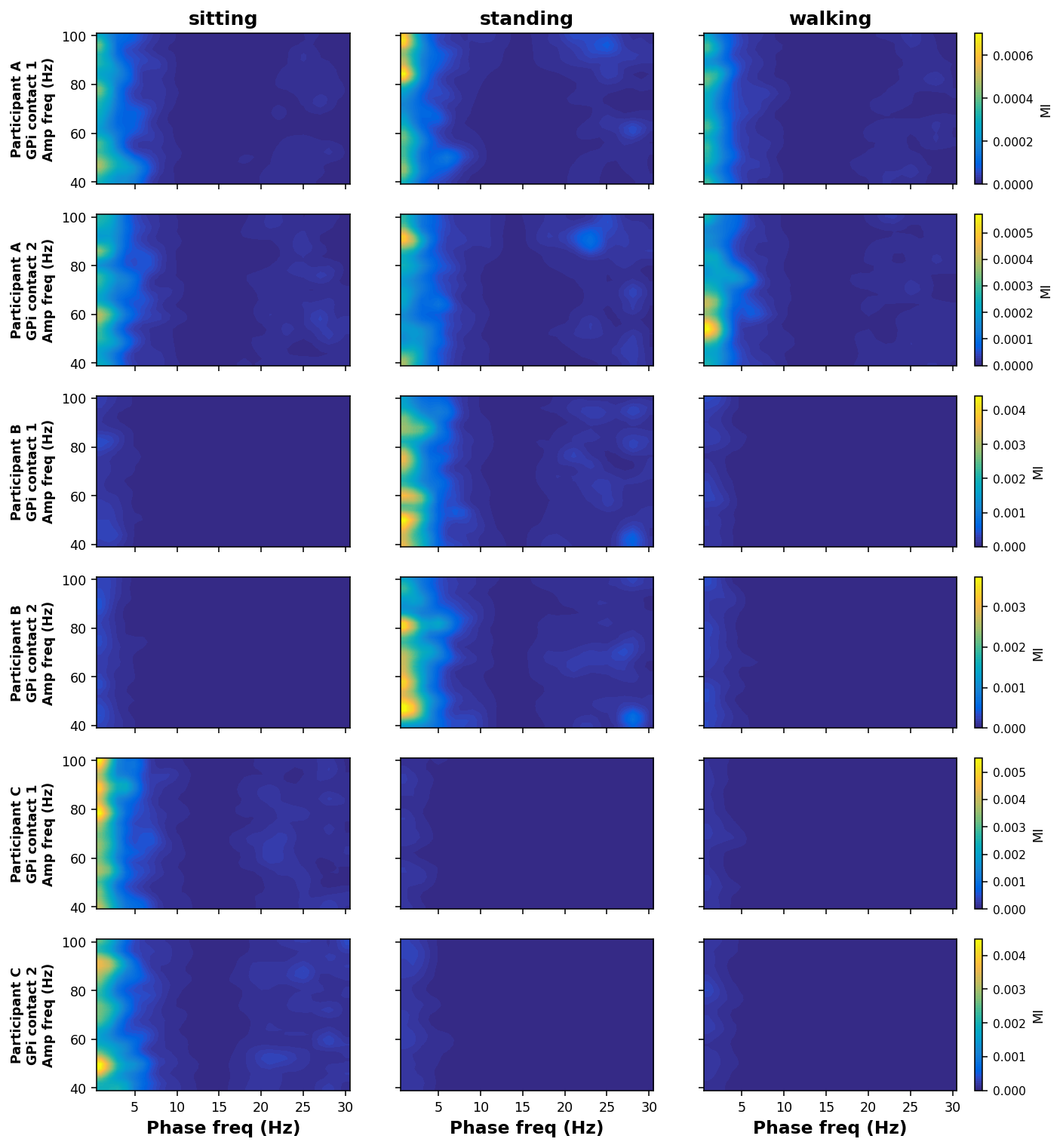

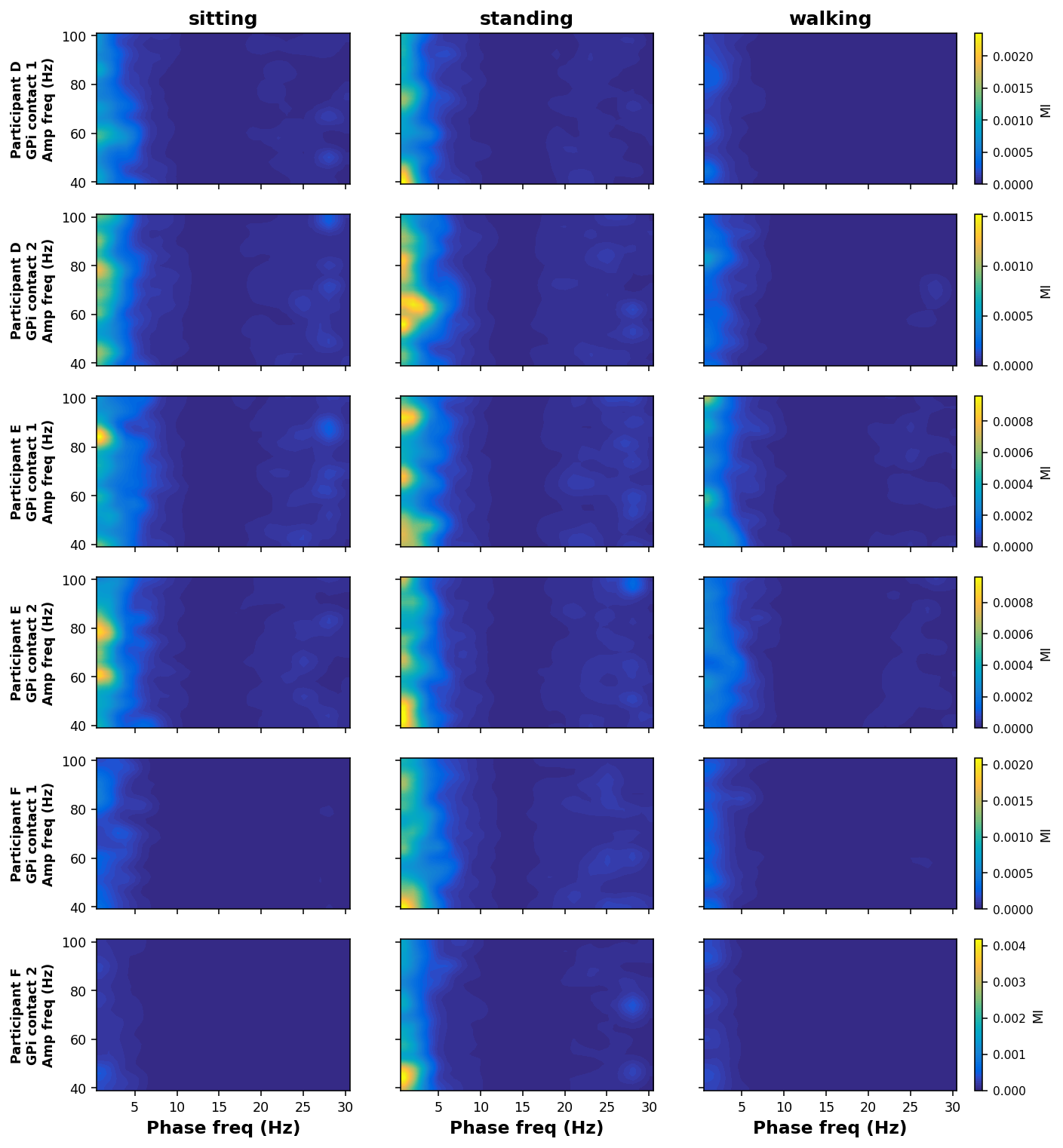

**Figure S3. Phase-Amplitude coupling comodulograms.** Each subplot represents the phase-amplitude comodulgrams for one of two GPi contacts used for each participant, across the three locomotor states. The three comodulograms within each subplot share the displayed color scale.

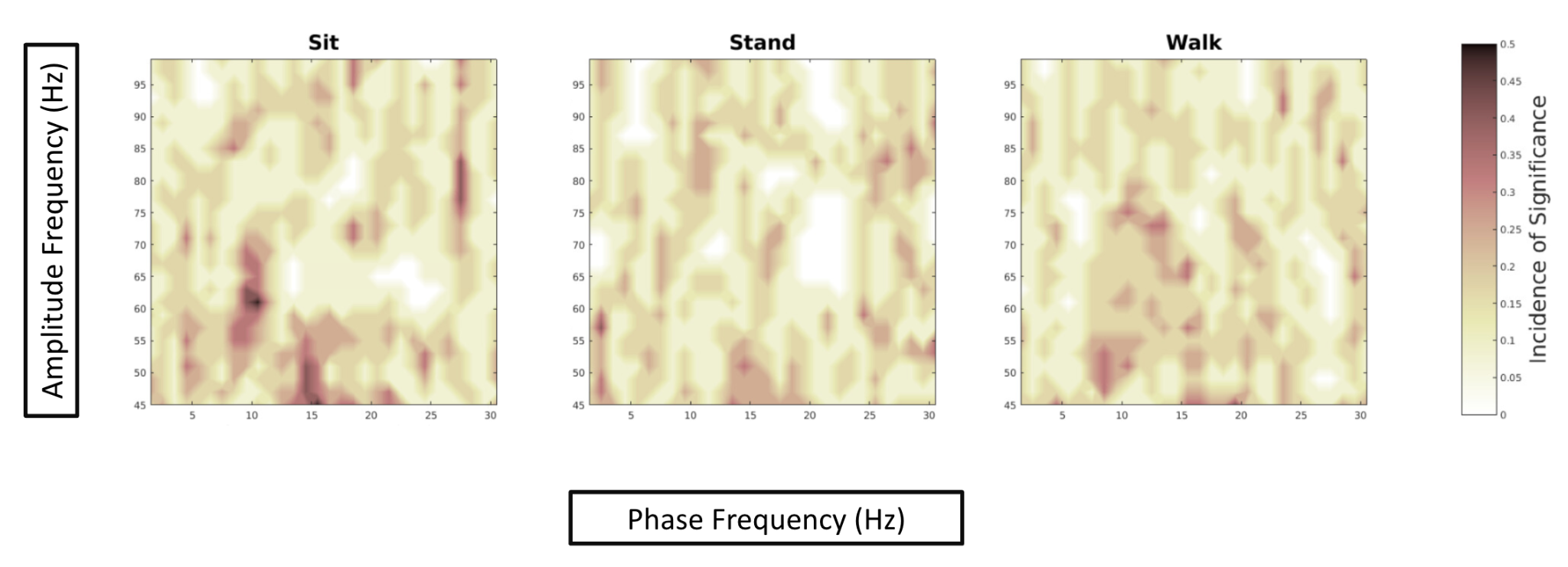

**Figure S4. Results of PAC surrogate testing.** Incidence of GPi contacts across all the participants with statistically significant phase-amplitude pairs. Significance was determined for a given phase-amplitude pair if the p-value for that pair was less than 0.05.

***Clinical Correlations***

|  | MDS-UPDRS-III | | nFoG-Q | |
| --- | --- | --- | --- | --- |
| Freq. Band | Sit -> Walk | Stand -> Walk | Sit -> Walk | Stand -> Walk |
| Delta | *Not tested* | R^2^ = 0.82  q = 0.04* | *Not tested* | R^2^ = 0.25  q = 0.47 |
| High Beta | R^2^ = 0.01  q = 0.93 | *Not tested* | R^2^ = 0.06  q = 0.65 | *Not tested* |
| Gamma | R^2^ = 0.00  q = 0.93 | *Not tested* | R^2^ = 0.35  q = 0.47 | *Not tested* |

**Figure S5A. Correlation between absolute change in bandpower between locomotor states and clinical scores using linear regression.** Bandpower modulation vs clinical scores. The three state-significant bandpower comparisons (BH-FDR family, m = 3) are reported for each clinical score, each at the state-pair that survived state-comparison screening. No outliers were flagged in any regression. * indicates q < 0.05. Grayed out boxes labeled “not tested” denote spectral measures that did not show statistical significance between locomotor states.

|  | UPDRS | | nFoG-Q | |
| --- | --- | --- | --- | --- |
| Freq. Band | Sit -> Walk | Stand -> Walk | Sit -> Walk | Stand -> Walk |
| Theta-gamma | R²=0.00, q=0.99 | R²=0.44, q=0.37 | R²=0.24, q=0.34 | n=6: R²=0.24, q=0.36  n=5: R²=0.97, q=0.006* |
| Alpha-gamma | R²=0.00, q=0.99 | R²=0.33, q=0.47 | R²=0.23, q=0.34 | n=6: R²=0.22, q=0.36  n=5: R²=0.98, q=0.006* |
| Beta-gamma | R²=0.00, q=0.99 | n=6: R²=0.43, q=0.43  n=5: R²=0.84, q=0.14 | R²=0.27, q=0.34 | n=6: R²=0.22, q=0.36  n=5: R²=0.96, q=0.007* |
| Low beta-gamma | R²=0.00, q=0.99 | R²=0.46, q=0.37 | R²=0.26, q=0.34 | n=6: R²=0.21, q=0.36  n=5: R²=0.93, q=0.015* |
| High beta-gamma | R²=0.00, q=0.99 | n= 6: R²=0.41, q=0.43  n=5: R²=0.84, q=0.14 | R²=0.29, q=0.34 | n=6: R²=0.22, q=0.36  n=5: R²=0.98, q=0.006* |

**Figure S5B. Correlation between absolute change in PAC between locomotor states and clinical scores using linear regression.** PAC modulation vs clinical scores across all 10 PAC band × walking-state-pair comparisons within each BH-FDR family (m = 10). Each cell shows R² and q for the full-cohort regression (n = 6) and, where an outlier was flagged by the Bonferroni-corrected outlier test on externally studentized residuals, the outlier-excluded regression (n = 5). Subject 0411 was flagged in all five Stand→Walk regressions against nFOG-Q; subject 0512 was flagged in β-γ and high β-γ Stand→Walk regressions against MDS-UPDRS-III. * indicates q < 0.05.
